# Presence of a home cage running wheel, but not wheel running per se, decreases social motivation in adult C57BL/6J female mice

**DOI:** 10.1101/2025.09.25.678626

**Authors:** Patryk Ziobro, Cassidy A. Malone, Stephen Batter, Longao Xu, Selina B. Xu, Alina Loginov, Katherine A. Tschida

## Abstract

Physical activity offers myriad benefits to health and well-being, in humans and other animals as well. In rodents, voluntary wheel running can attenuate the effects of both physical and social stressors on rodent social behavior. Whether wheel running affects rodent social behaviors per se remains less well understood. We conducted the current study to test whether home cage access to running wheels impacts the social behaviors of adult, group-housed C57BL/6J female mice during same-sex interactions with novel females. Group-housed females were either given continuous home cage running wheel access or a standard paper hut starting at weaning, and as adults, social behaviors were measured during interactions with novel females. In two cohorts, we found that 5 weeks of running wheel access during adolescence reduced the time that subject females spent investigating a novel female and also tended to reduce total ultrasonic vocalizations produced during interactions. These effects were not reversed by a 2-week period of running wheel removal but were recapitulated in a different cohort by 2 weeks of running wheel access in adulthood. Unexpectedly, we found that these effects on female social behavior were not due to wheel running per se, because females raised from weaning with ‘immobile’ running wheels also showed low rates of social behaviors during same-sex interactions in adulthood. Overall, we find that the presence of a running wheel in the home cage has an enduring inhibitory influence on female social behavior during same-sex interactions, a finding that has implications for the design of studies that include same-sex interactions between female mice.

## Introduction

Physical activity (i.e., exercise) promotes both physical and mental well-being. For example, exercise reduces the risk of chronic diseases, including cardiovascular disease, diabetes, and cancer (1–8). Exercise is also associated with improved cognitive function (9,10), reduced depressive symptoms (4,11–15), and better health-related quality of life (16–18).

Similar to findings in humans, studies in rodents also support the idea that exercise conveys health benefits, including cardiovascular benefits (19,20), increased cognitive ability and hippocampal neurogenesis (21,22), and antidepressant and anxiolytic effects on behavior (19,23– 25). The most common form of exercise provided to laboratory rodents is voluntary wheel running, in which rodents are given free access to a running wheel in their home environment (20,26). In fact, both laboratory and wild rodents run spontaneously when given access to running wheels (20,26,27), indicating that rodents find wheel running to be rewarding (28–30). Within the vast literature examining the effects of voluntary wheel running on rodent health and behavior, there is a particularly rich body of work showing that wheel running can ameliorate stress-induced changes in social behavior. For example, wheel running attenuates the effects of chronic social defeat stress on sociability and aggression in male mice (31–33), attenuates the effects on sociability of inescapable tail shock in female and male rats (34,35), reduces the effects of chronic social isolation on sociability and aggression in male mice (36), and reduces the severity of anxiety-like and depressive-like behaviors in female prairie voles subjected to a combination of social isolation and chronic mild stress (37).

Although past work robustly supports the idea that voluntary wheel running can attenuate the effects of stressors on rodent social behavior, much less is known about whether wheel running impacts rodent social behaviors per se. This gap in knowledge stems from the fact that studies employing voluntary wheel running typically include single-housed subjects with free access to a running wheel that electronically tracks rotations, in order to longitudinally track wheel running for a single animal. Social isolation itself is a stressor that can substantially alter mouse social behaviors (38–43), which complicates the use of the standard voluntary wheel running paradigm to investigate the effects of wheel running on social behavior. Thus, whether wheel running affects the social behaviors of unstressed, socially-housed rodents remains understudied.

To provide mice with exercise enrichment, our lab routinely places (non-electronic) running wheels in mouse home cages at the time of weaning, and cages of mice in our colony subsequently have continuous running wheel access throughout adolescence and adulthood. Given our lab’s interest in the behavioral and social contexts that influence affiliative social behaviors of female mice (42–44), we designed the current study to test whether free access to running wheels impacts the social behaviors of adult, group-housed females. Because we provided groups of females with non-electronic running wheels, we use the term “running wheel access” to distinguish our experimental design from the standard voluntary wheel running paradigm. In Experiment 1, same-sex groups of female siblings were either given continuous home cage running wheel access or a standard paper hut starting at weaning (3 weeks of age), and as adults (8 weeks of age), the social behaviors of females from these two groups were measured during interactions with novel female mice. In follow-up experiments, we replicated our findings from Experiment 1 in a second cohort of females (Experiment 2a), tested whether the effects of 5 weeks of running wheel access on female social behavior could be reversed by a 2-week period of running wheel removal (Experiment 2b), and tested whether a 2-week period of running wheel access in adulthood could recapitulate the effects of 5 weeks of running wheel access throughout adolescence on female adult social behaviors (Experiment 2c). Finally, to understand whether impacts on female social behaviors were due to wheel running per se, we compared adult social behaviors in females that were raised from weaning with either intact, mobile running wheels or altered, ‘immobile’ running wheels. Our findings suggest that the presence of a home cage running wheel exerts a generally inhibitory effect on female social behavior, but that these effects are not due to wheel running per se.

## Materials and Methods

Further information and requests for resources should be directed to the corresponding author, Katherine Tschida.

### Ethics Statement

All experiments and procedures were conducted according to protocols approved by the Cornell University Institutional Animal Care and Use Committee (protocol #2020-001).

### Subjects

Adult (> 8 weeks old) female C57BL/6J (Jackson Laboratories, 000664) mice were group-housed with same-sex siblings from weaning (postnatal day 21) until the start of experiments. In Experiments 1 and 2, weanling females were assigned to either the hut condition (control) or the wheel condition (experimental). Hut condition cages included a paper hut and other standard enrichment (crinkle paper, nestlets, and care fresh bedding) but no running wheel (Fig. 1, left). Wheel condition cages included most standard enrichment (crinkle paper, nestlets, and care fresh bedding) but did not contain a paper hut and instead contained a combined shelter and running wheel (Innodome + Innowheel, from Innovive) (Fig. 1, right). In Experiment 3, weanling females were assigned to either the mobile wheel condition or the immobile wheel condition (see below), and cages for both of these groups included standard all enrichment except for a paper hut. Mice were kept on a 12h:12h reversed light/dark cycle and given ad libitum food and water for the duration of the experiment. Subjects did not have social interactions with novel mice prior to the start of experiments.

**Figure 1.**
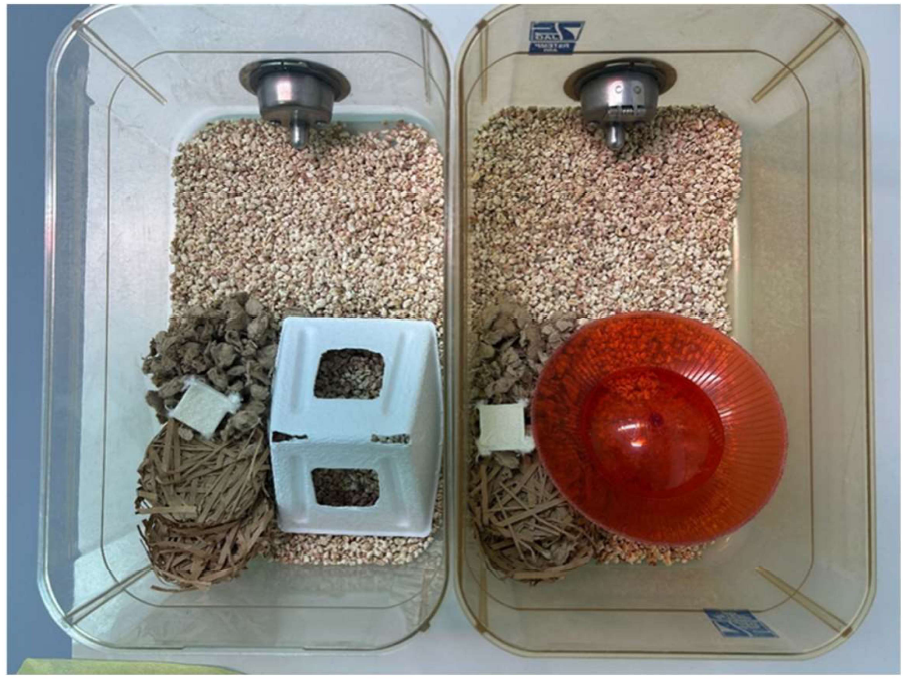
Home cage contents for cages in hut vs. wheel conditions. Home cage contents are shown for a cage in the hut condition (left) and a cage in the wheel condition (right). Food and water were provided at libitum but are not shown.

### Experiment 1 design (Fig. 2, top)

**Figure 2.**
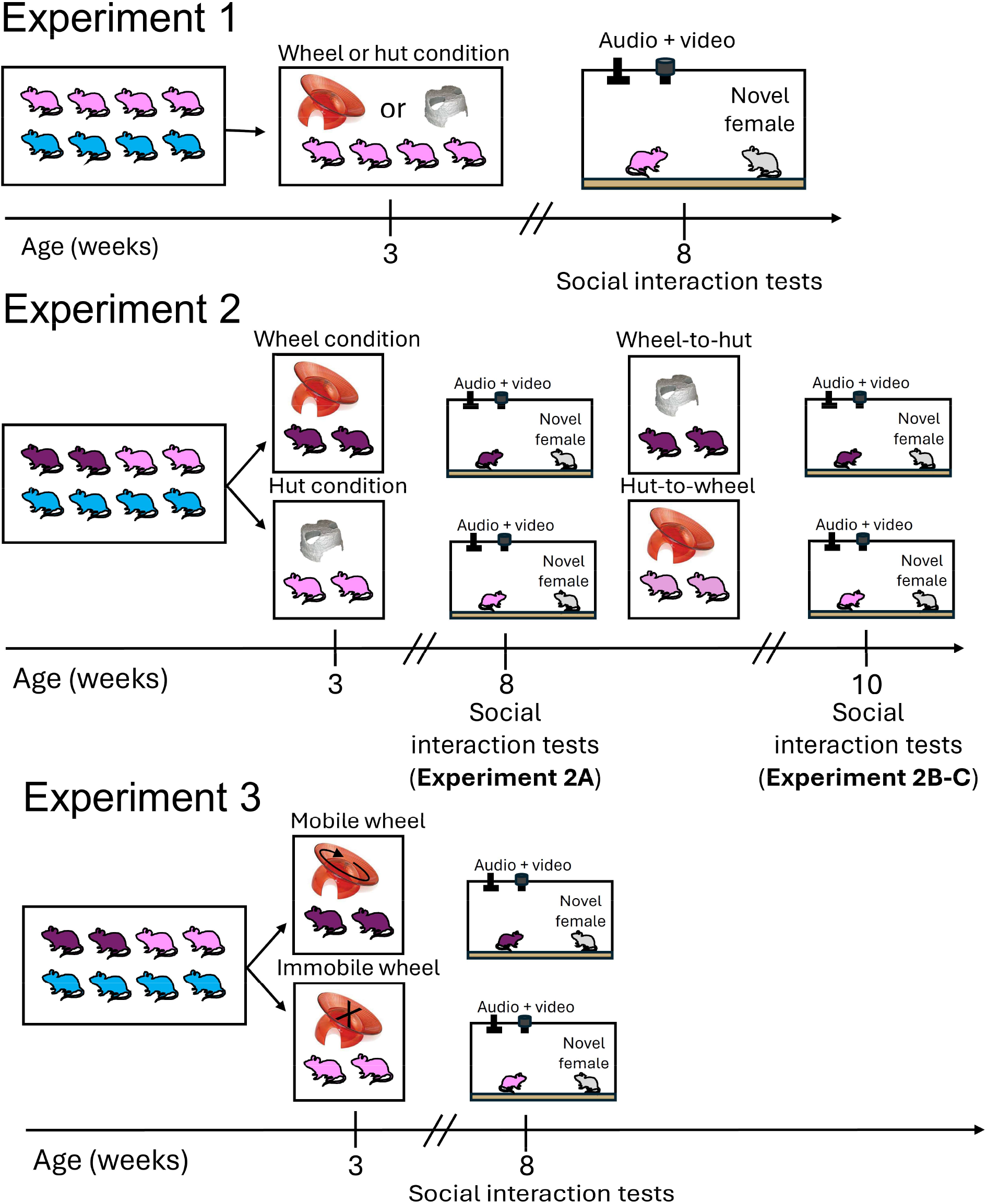
Study design. Schematic of the study design is shown for Experiment 1 (top), Experiment 2 (middle), and Experiment 3 (bottom). Female subjects are shown in light and/or dark purple, male siblings are shown in blue, and novel females used in social interaction tests are shown in gray.

At the time of weaning (postnatal day 21), female siblings from a given litter were placed into a home cage and housed together until adulthood. In Experiment 1, each female sibling group was assigned to either the wheel or the hut condition, and cages were maintained in their respective enrichment conditions until adulthood. All cages assigned to the wheel condition had continuous access to a running wheel, but running wheel usage by individual mice within the cage was not tracked. After 5 weeks in either the wheel or the hut condition (∼8 weeks of age), subject females were given a social interaction with a novel, group-housed female mouse. On the day of the social interaction test, the home cage of the subject female was placed into a sound-attenuating recording chamber (Med Associates) and siblings were moved temporarily to a clean cage. A custom lid (with tall sides and no top) was placed on the cage to allow video and audio recordings, and the mouse shelter (either running wheel or paper hut) was removed from the cage. The recording chamber was equipped with an infrared light source (Tendelux) and a webcam (Logitech), with the infrared filter removed to enable video recording under infrared lighting. A novel, group-housed adult female was then placed in the cage with the subject and allowed to interact for 10 minutes. Novel females were re-used across multiple social interaction tests, but the same novel female was never used in more than one social interaction test per day.

### Experiment 2 design (Fig. 2, bottom)

The goals of Experiment 2 were two-fold. First, we aimed to replicate the findings of Experiment 1 and to account for potential contributions of litter effects to the results of Experiment 1. To this end, Experiment 2 employed a design in which females from a given litter were divided into two groups at weaning, with one group of female siblings placed in a home cage and assigned to the wheel condition, and the remaining female siblings placed in another home cage and assigned to the hut condition. As a result, only litters that included at least 4 female siblings were used in Experiment 2, such that each female cage contained 2-3 siblings. Second, we wanted to test whether removing running wheels from cages of adult females from the wheel condition would reverse the effects of 5 weeks of running wheel access on female social behavior. Conversely, we wanted to evaluate whether a shorter period of running wheel access in adulthood could mimic the effects on female social behavior of 5 weeks of running wheel access from weaning until adulthood. To this end, Experiment 2 females were assigned to the wheel or hut condition at weaning and then given a 10-minute-long social interaction with a novel, group-housed female at 8 weeks of age as described for Experiment 1. Then, each female cage’s condition was reversed for 2 weeks (i.e., running wheels were removed from wheel condition cages and replaced with paper huts, and paper huts were removed from hut condition cages and replaced with running wheels), and female subjects were given a second 10-minute-long social interaction test at 10 weeks of age. Social interaction tests at 8 weeks and 10 weeks were conducted as described in Experiment 1, and each subject female was tested with a different (novel) female at the 8-week and 10-week social interaction tests.

### Experiment 3 design (Fig. 2, bottom)

Experiment 3 also employed a design in which females from a given litter were divided into two groups at weaning, with one group of female siblings assigned to the mobile wheel condition, and the remaining female siblings assigned to the immobile wheel condition. Running wheels were made immobile by using a heat gun to soften the plastic of the running wheel base at its connection point with the bottom tab of the running wheel and then using pliers to squeeze and deform the connection, rendering the running wheel unable to rotate but otherwise identical to the mobile running wheels. After 5 weeks in either the mobile wheel or the immobile wheel condition (∼8 weeks of age), subject females were given a 10-minute social interaction with a novel, group-housed female mouse as described above.

### Scoring of social investigation from video recordings

Trained observers blinded to experimental group assignments used BORIS v.8.13 to score times in webcam videos during which the subject female engaged in social investigation (i.e., sniffing or following) of the novel female. Times marked by mutual social investigation (mutual sniffing and/or circling) were also included in the total amount of social investigation time for the subject female. We did not observe any instances of mounting or fighting in our dataset.

### USV recording and detection

USVs were recorded with an ultrasonic microphone (Avisoft, CMPA/CM16), amplified (Presonus TubePreV2), and digitized at 250 kHz (Spike 7, CED). USVs were detected with custom Matlab codes (Tschida et al., 2019) using the following parameters: mean frequency > 45 KHz; spectral purity > 0.3; spectral discontinuity < 1.0; minimum USV duration = 5 ms; minimum inter-syllable interval = 30 ms). Because we did not use methods to assign USVs to individual signalers (44–48), total USVs reported in social interaction tests represent the total number of USVs produced by each pair of females.

### Statistical analyses

The Shapiro-Wilk test was used to evaluate the normality of data distributions, and non-parametric two-sided comparisons were used for non-normally distributed data (alpha = 0.05). No statistical methods were used to pre-determine sample size.

## Results

### Experiment 1: Running wheel access throughout adolescence decreases social investigation in adult C57BL/6J female mice during interactions with novel females

After 5 weeks of continuous home cage access to either a running wheel or a paper hut, each group-housed adult (8-week-old) female subject was given a 10-minute-long social interaction test with a novel, group-housed female mouse (Fig. 2; see Methods). Given prior literature on the benefits of voluntary wheel running in rodents, we hypothesized that long-term running wheel access would promote social motivation in adult female mice and in turn would enhance rates of social investigation and ultrasonic vocalizations (USVs) during affiliative social interactions with novel females. Contrary to our hypothesis, we found that female subjects from the wheel condition spent significantly less time engaged in social investigation of novel, visitor females compared to female subjects in the hut condition (Fig. 3A, left; Mann Whitney U test: z = 3.1, p = 0.002). Total USVs also tended to be lower in pairs that included females from the wheel condition, although this difference was not statistically significant (Fig. 3A, right; Mann Whitney U test: z = 1.9, p = 0.06). In summary, running wheel access throughout adolescence decreased social investigation in adult female mice during interactions with novel female mice and also tended to decrease total USVs produced by the pair.

**Figure 3.**
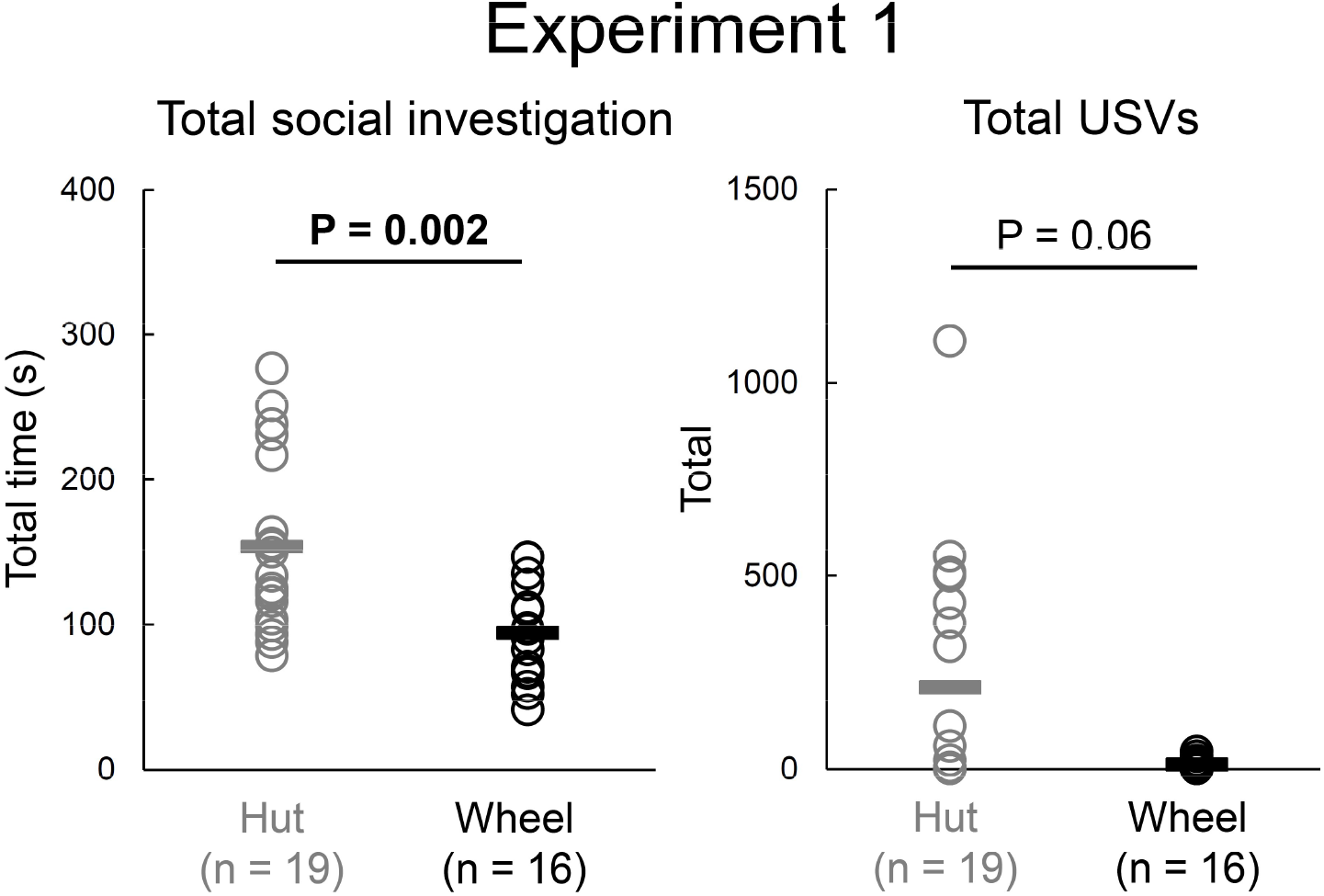
Running wheel access throughout adolescence decreases social investigation in adult female mice during interactions with novel females. (A) Left plot shows total time that female subjects spent engaged in social investigation of a novel female. Right, same as left, for total USVs recorded from female-female pairs. Black and gray lines show mean values.

### Experiment 2a: In a second cohort of adult C57BL/6J females, running wheel access throughout adolescence decreases social investigation and total USVs during interactions with novel females

One caveat to the study design in Experiment 1 is that all female siblings from a given litter were assigned to either the wheel condition or to the hut condition. Thus, differences in social behavior between litters might have contributed to the differences in social behavior between females in the wheel and hut conditions. To better account for any contribution of litter effects, we conducted a second set of experiments with a modified design, in which female siblings from each litter were divided, with a subset (≥ 2 female siblings) assigned to the wheel condition and the remaining female siblings (≥ 2) assigned to the hut condition (Fig. 2, middle).

Similar to the results of Experiment 1, we found that females from the wheel condition spent significantly less time engaged in social investigation of novel females compared to females in the hut condition (Fig. 4, left; Mann Whitney U test: z = -4.45, p < 0.0001). Total USVs were also significantly lower in pairs that included females from the wheel condition (Fig. 4, right; Mann Whitney U test: z = -2.34, p = 0.02). Alongside the results of Experiment 1, these findings support the idea that running wheel access throughout adolescence decreases social motivation in adult female mice during same-sex interactions.

**Figure 4.**
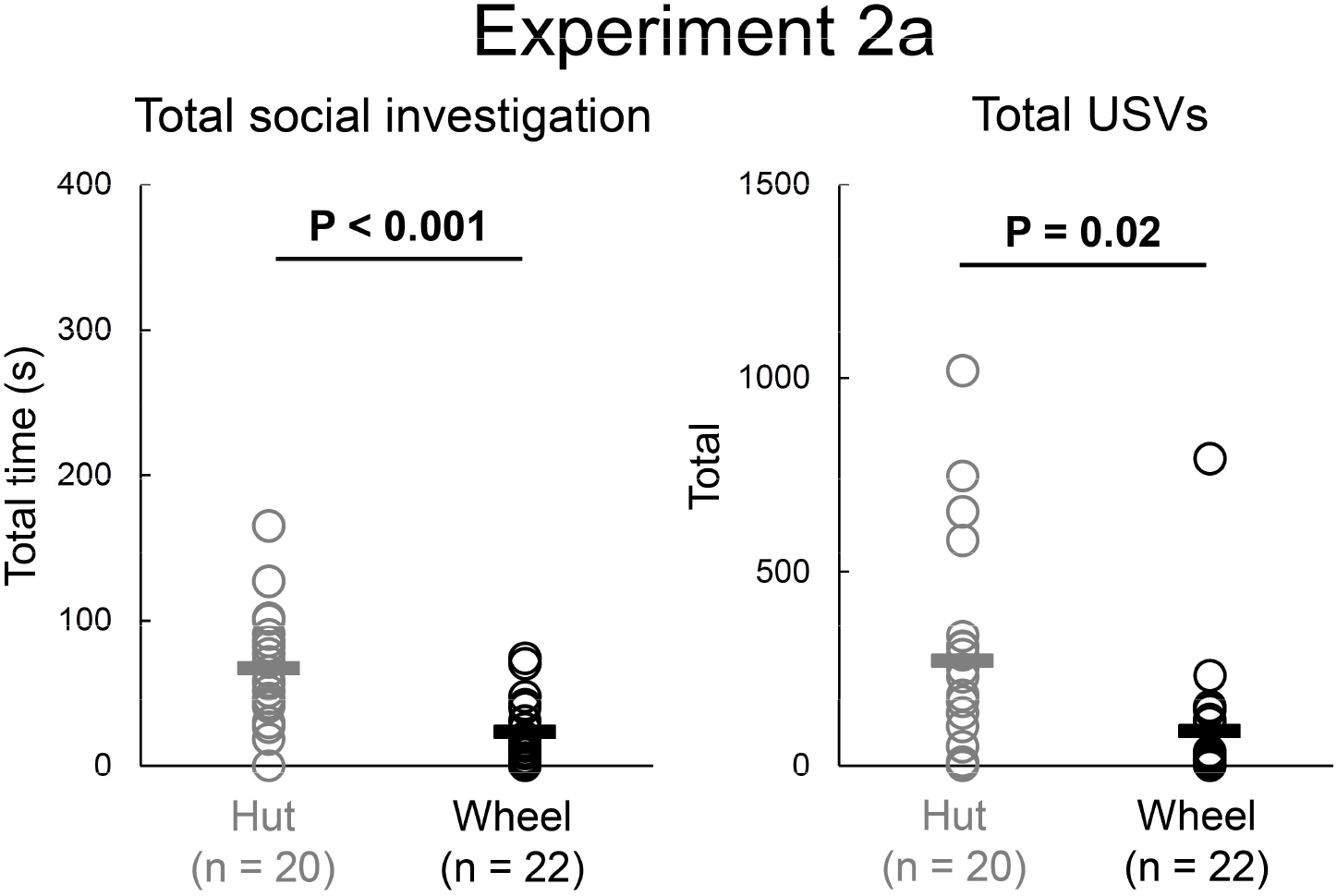
In a second cohort of females, running wheel access throughout adolescence decreases social investigation and total USVs during interactions with novel females. Left plot shows total time that female subjects spent engaged in social investigation of a novel female. Right, same as left, for total USVs recorded from female-female pairs. Black and gray lines show mean values.

### Experiment 2b: Can removal of running wheels in adulthood reverse the effects of running wheel access throughout adolescence on adult female social behavior?

Given the robust effects of long-term running wheel access on the social behaviors of adult female mice, we next tested whether these effects could be reversed by removing the running wheel. To this end, after their 8-week social interaction tests, females (n = 22) from Experiment 2a that had been raised with a running wheel since weaning had the running wheels removed from their home cages and replaced with paper huts. After 2 weeks (10-week time point), the wheel-to-hut females were given a second social interaction test with a novel female (Fig. 2, middle). We found that removal of running wheels for 2 weeks failed to significantly increase either social investigation or total USVs (Fig. 5, top; Wilcoxon signed-rank tests to compare 8-week vs. 10-week behaviors; z = -1.18, p = 0.24 for social interaction; z = -1.34, p = 0.18 for USVs).

**Figure 5.**
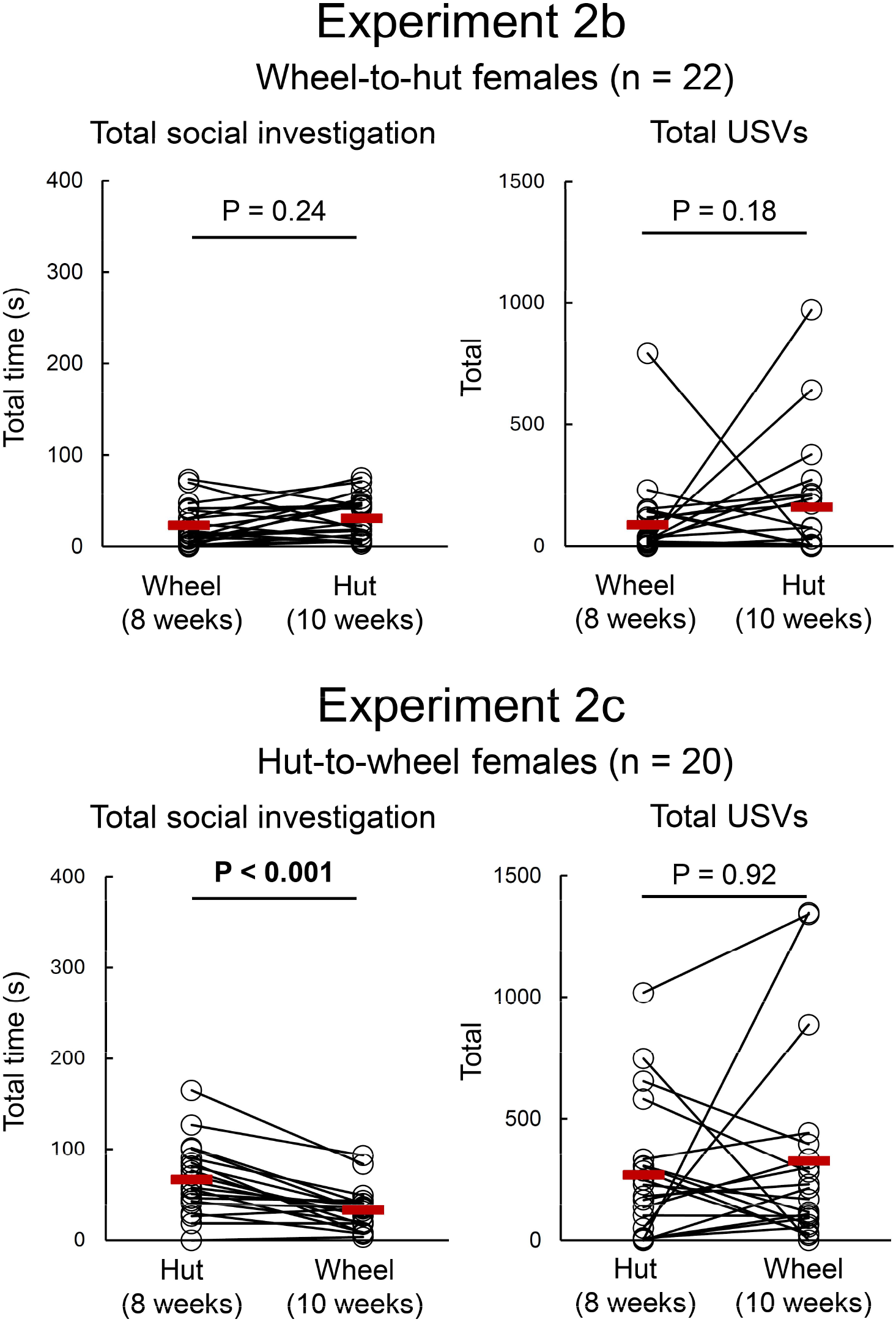
Effects of adulthood removal of running wheels or adulthood-only running wheel access on female social behavior. (Top) Left plot shows total time that female subjects spent engaged in social investigation of a novel female, after 5 weeks of running wheel access (8-week time point) and again after 2 weeks with the running wheel removed and replaced with a paper hut (10-week time point). Right, same as left, for total USVs. Red symbols show mean values. (Bottom) Left plot shows total time that female subjects spent engaged in social investigation with a novel female after 5 weeks of paper hut access (8-week time point) and again after 2 weeks of running wheel access (10-week time point). Right, same as left, for total USVs. Red symbols show mean values.

### Experiment 2c: Can 2-weeks of running wheel access in adulthood mimic the effects of running wheel access throughout adolescence on adult female social behavior?

Multiple factors could contribute to the inhibitory effects of running wheel access on adult female social behavior observed in Experiments 1 and 2a. One possibility is that the timing of running wheel access during adolescence is a critical factor. Another possibility is that the total duration (∼5 weeks) of running wheel access is important, and that shorter durations of running wheel access, regardless of when they occurred during development, would fail to impact female social behavior. On the other hand, it could be that even a shorter duration of running wheel access in adulthood is enough to attenuate female social behavior during interactions with novel females. To test this last idea, after their 8-week social interaction test, females (n = 20) from the hut condition in Experiment 2a had their paper huts removed and replaced with running wheels. After 2 weeks (at 10-weeks of age), the hut-to-wheel females were given a second social interaction test with a novel female (Fig. 2, middle).

Two weeks of running wheel access significantly decreased the total time that females engaged in social investigation of a novel female (Fig 5, bottom left; Wilcoxon signed-rank tests to compare 8-week vs. 10-week total social investigation; z = -3.57, p < 0.001). Notably, total social investigation in the hut-to-wheel females (after 2 weeks of running wheel access in adulthood) was not significantly different from total social investigation in the Experiment 2A females that had 5 weeks of running wheel access throughout adolescence (Mann Whitney U test: z = -1.62, p = 0.10). In contrast to the decrease in social investigation, there was no significant change in total USVs recorded from pairs that included the hut-to-wheel females (Fig. 5, bottom right; Wilcoxon signed-rank tests to compare 8-week vs. 10-week total USVs; z = -0.1, p = 0.92). We conclude that at least with regard to time spent engaged in social investigation, the effects of longer-term running wheel access during adolescence can be recapitulated by a shorter period of running wheel access in adulthood.

### Experiment 3: Is wheel running per se required for the effects of home cage running wheels on adult female social behavior?

The results of Experiments 1 and 2 suggest that exercise (wheel running) may exert an overall inhibitory effect on female social behaviors. However, running wheels and paper huts differ in multiple ways, both in their physical features as well as how mice can interact with them. To test whether running per se accounts for the effects of home cage running wheel access on female social behavior, we set up a final cohort of females in which female siblings from each litter were divided and assigned to either the ‘mobile’ running wheel condition (identical to the running wheel condition in Experiments 1-2) or to the ‘immobile’ running wheel condition, which included a running wheel that was altered to prevent it from rotating but was otherwise identical to the mobile running wheels (see Methods) (Fig. 3). At 8 weeks of age, females from both groups were given social interaction tests with novel females.

Unexpectedly, we found that females from the immobile and mobile wheel conditions did not differ in the time they spent engaged in social investigation of novel females (Fig. 6, left; Mann Whitney U test: z = 0.24, p = 0.81), nor did the groups differ in total USVs (Fig. 6, right; Mann Whitney U test: z =1.01, p = 0.31). Moreover, both of the Experiment 3 groups were significantly lower in their social investigation times than the Experiment 2 females raised with huts but were not different from the Experiment 2 females raised with (mobile) wheels (X^2^(3) = 16.5, p = < 0.001; p < 0.05 for Experiment 2 hut females vs. all other Experiment 2 and 3 groups; Kruskal-Wallis test with Dunn’s post-hoc test). These findings suggest that presence of home cage running wheels, but not wheel running per se, decreases social motivation in adult female mice during same-sex interactions.

**Figure 6.**
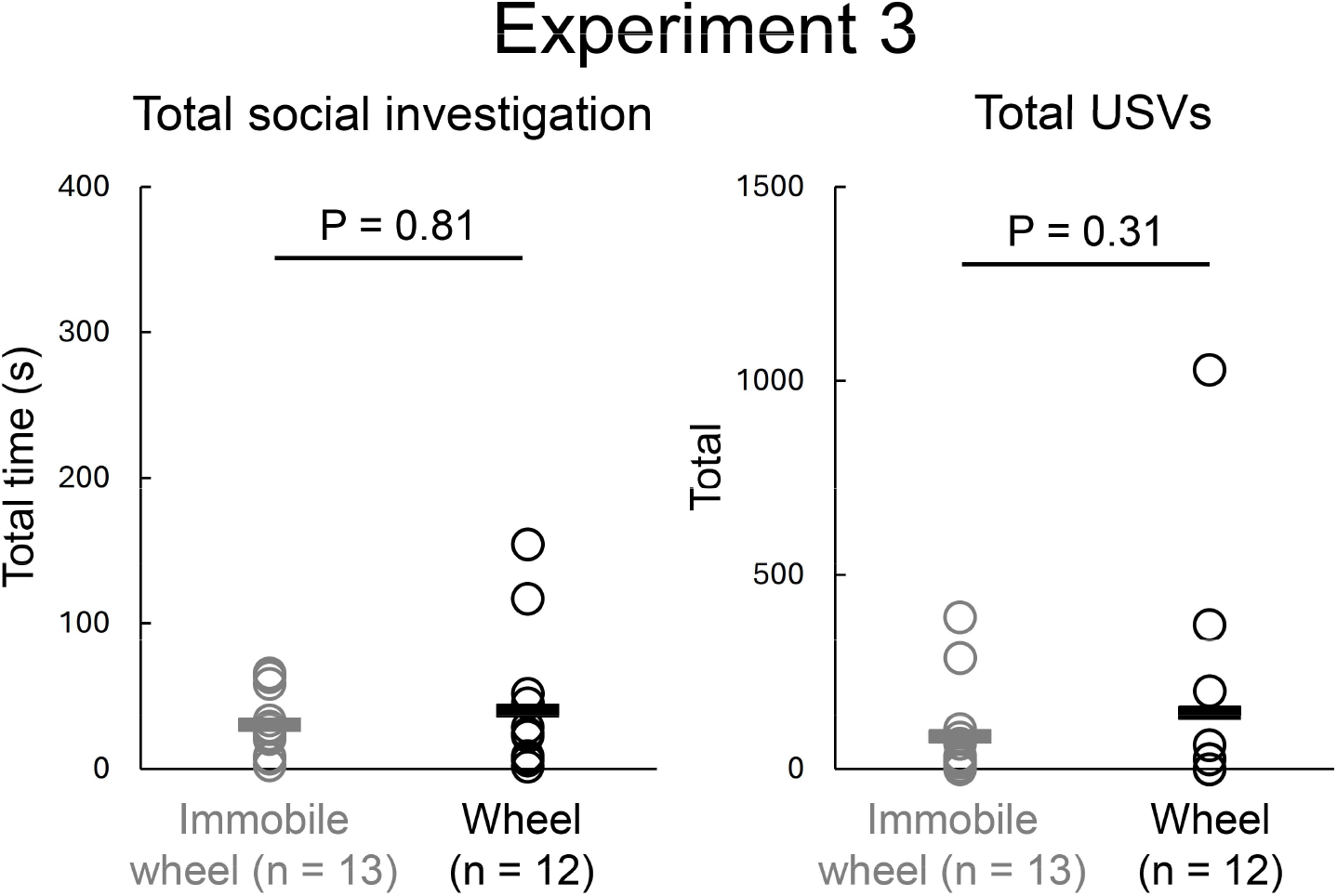
Females raised with mobile and immobile running wheels do not differ in their social investigation time and total USVs during interactions with novel females. Left plot shows total time that female subjects spent engaged in social investigation of a novel female. Right, same as left, for total USVs recorded from female-female pairs. Black and gray lines show mean values.

## Discussion

Across two cohorts of female mice, we find evidence that 5 weeks of running wheel access throughout adolescence decreases social motivation in adulthood during interactions with novel females. These effects were not reversed by a 2-week period of running wheel removal, but the finding that 2 weeks of running wheel access in adulthood also reduces female social investigation suggests that these effects are not specific to early-life development. Unexpectedly, we found that our results cannot be attributed due to wheel running per se, because females that were raised with immobile running wheels also displayed low rates of social investigation and USV production during same-sex interactions. What mechanisms can account for these inhibitory effects of home cage running wheels on female social behavior?

One possibility is that differences in the physical features of running wheels and paper huts drive changes in female social behavior. The running wheels we used in the current study are semi-transparent to light, and it is possible that reduced access to a dark location within the home cage could influence female social behavior. Another difference is that the running wheels are constructed of durable plastic and cannot be chewed to create additional nesting material, although we note that all cages in the wheel condition were provided with ample enrichment (crinkle paper, nestlets, and care fresh bedding). An additional possibility is that the presence of running wheels, whether mobile or immobile, drives some difference in home cage behaviors that accounts for our findings. For example, even though the running wheels include a base that mice can use as a nesting location, we have observed anecdotally that cages with running wheels often create nests outside of the wheel base. Such nests may be more exposed than nests constructed under a standard paper hut, potentially leading to changes in anxiety, thermoregulatory behaviors, or other aspects of home cage behavior. More work is required to understand which running wheel characteristics drive the effects on female social behavior and whether these effects would be observed with other types of home cage running wheels.

The finding from Experiment 3 that mobile and immobile females do not differ in their social investigation times or total USVs suggests that at least in a standard home cage, wheel running does not promote female social behaviors as we had initially hypothesized. Our findings do not rule out the possibility that wheel running could promote affiliative social behaviors, perhaps in larger home cage environments that would enable the inclusion of a running wheel as well as paper huts or other potential nesting locations.

In summary, our results show that the presence of a home cage running wheel can exert unanticipated and long-lasting consequences on the social behaviors of female mice. Although there is ample evidence that wheel running offers important welfare benefits for laboratory rodents, our findings nonetheless have important implications for the design of studies that include measures of social behavior in female mice. Whether the presence of a home cage running wheel exerts similar effects on the social behaviors of male mice, either during same-sex interactions with other males or during courtship interactions with females, could also be explored in future studies.

## Acknowledgements

We thank the CARE staff for their excellent mouse husbandry. Thanks also to our reviewers for their insightful suggestion to conduct the immobile running wheel control experiments.

